# Treated eave screens in combination with screened doors and windows, are more effective than untreated eave screens in a similar combination in reducing indoor and outdoor *Anopheles* populations under semi-filed conditions in western Kenya

**DOI:** 10.1101/2023.07.05.547882

**Authors:** Bernard Abong’o, Silas Agumba, Vincent Moshi, Jacob Simwero, Jane Otima, Eric Ochomo

**Author notes:** Corresponding author –.

## Abstract

**Introduction:** Human habitats remain the main point of human-vector interaction leading to malaria transmission despite sustained use of insecticide treated nets (ITNs) and indoor residual spraying (IRS). Simple structural modifications involving screening of doors, windows and eaves have great potential for reducing indoor entry of mosquitoes and hence malaria transmission.

**Methods:** Four huts, each constructed inside a semi-field structure, allowing the experimental release of mosquitoes at the Kenya Medical Research Institute (KEMRI), Kisumu research station were used in the study. Two huts had untreated eave and door screens and screened air cavities in place of windows in experiment 1 with the eave screen treated using Actellic® insecticide in experiment 2. The other two huts remained unscreened throughout the study. First filial (F1) generation of *Anopheles funestus* from Siaya, F0 reared from *An. arabiensis* larvae collected from Ahero and *An. arabiensis* Dongola strain from the insectary were raised to 3-day old adults and used in experiments. Two hundred, 3-day old adults of each species were released in each semi-field structure at dusk and recaptured the following day at 0700hrs and at 0900 hours. A single volunteer slept in each hut under untreated bed net each night of the study. Recaptured mosquitoes were counted and recorded by collection location, either indoor or outdoor of each hut in the different semi-field structures.

**Results:** Significantly fewer *An. arabiensis* from Ahero [RR=0.10; (95%CI: 0.02-0.63); P<0.0145], *An. arabiensis* Dongola strain [RR=0.11; (95%CI: 0.04 – 0.19); P<0.0001 and *An. funestus* from Siaya [RR=0.10; (95%CI: 0.06-0.17); P<0.0001] were observed inside modified huts compared to unmodified ones. Treating of eave screen material with Actellic® 300CS significantly reduced the numbers *An. arabiensis* from Ahero [RR=0.003; (95%CI: 0.00-0.03); P<0.0001] and *An. arabiensis* Dongola strain [RR=0.03; (95%CI: 0.02-0.05); P<0.0001] indoors of huts with treated eave screen compared to huts with untreated eave screens, while totally preventing entry of *An. funestus* indoors. These modifications cost <250usd/structure.

**Discussion and Conclusion:** This article describes affordable and effective ways of reducing mosquito entry into the house by modifying the eaves, doors and windows. These modifications were highly effective in reducing indoor entry of mosquitoes. Additionally, treatment of eave screen material with an effective insecticide further reduces the *Anopheles* population in and around the screened huts under semi-field conditions and could greatly complement existing vector control efforts.

## Introduction

The world recently reported an increase in estimated malaria cases in the year 2021 compared to 2020 [1]. Most of this increase in malaria cases and the greatest disease burden were reported to come from Sub-Saharan Africa region where malaria control is heavily reliant on use of insecticidal treated nets (ITNs), artemisinin-based combination therapy (ACT) and indoor residual spraying (IRS). ITNs was observed to have contribute the greatest to decline in malaria transmission between the year 2000 and 2015 [2]. However, more recently, little or no progress in reduction of malaria cases has been witnessed despite sustained use of these interventions [1, 3].

Both ITNs and IRS are insecticide based, malaria vector control interventions whose implementations are limited to personal protection at bedtime and application on the walls respectively. A number of limitations including insecticide resistance in mosquitoes [4-7], incomplete coverage [8-10], low compliance to ITN use [11-13] and changing vector behaviour [14-17] are potentially lessening the effectiveness of these interventions. Consequently, complementary mosquito control tools are urgently required to sustain the gains made in malaria control and to further reduce the disease burden.

Malaria transmitting mosquitoes are closely associated with human habitation. Mosquitoes are known to enter and bite within human houses at night [18], thereby transmitting malaria. Even though reports of changing vector biting behaviour due to sustained ITN use exist [4, 19-21], the greatest bulk of malaria transmission still occurs indoors in many malaria endemic settings [22] (Ichodo *et al* in prep). In western Kenya, recent studies have reported persistently, high, late night biting malaria vectors [23-25]. Consequently, to control malaria transmission in these regions, it is important to first identify the location of human-vector interaction to effectively targe control efforts. The alteration of house designs to limit indoor mosquito entry offers unparalleled and non-insecticide-based potential that could effectively complement malaria vector control efforts over a longer period.

Housing is a key contributor to health; it not only protects against the elements but also influences the physical and psychosocial well-being of its inhabitants. House modification has been reported in many settings to be an effective mosquito control strategy [26-29] including resulting in an epidemiological impact [27]. However, it has received little attention in much of Sub-Saharan Africa where malaria is endemic likely due to the lack of a standardized approach and perceived cost. The World Health Organization (WHO) Global Malaria Programme recently provided a conditional recommendation for house modification based on low certainty of evidence [30]. The impact of screening doors, windows and eaves, followed by the treatment of eave screen material with an effective insecticide was evaluated in a semi field condition to provide evidence for improved housing as a malaria control intervention and practical lessons for community implementation of a sustainable vector control intervention in western Kenya.

### Methodology

The experiments were conducted at the Kenya Medical Research Institute – Centre for Global Health Research (KEMRI – CGHR) Kisumu. The campus has four semi-field stations which are double netted, double door structures measuring 20m length and 8m wide and rise to 4.5m at the apex [31, 32]. We modified the semi-field structures with a 3m high netting to ensure ease of mosquito recapture. Each semi-field structure houses one hut measuring 3m long X 3m wide X 2m high. The huts have an open ceiling and wooden doors and windows (1 each) (Figure 2). These huts are similar to a typical simple house structure in the western Kenya region (Figure 1).

**Figure 1:**
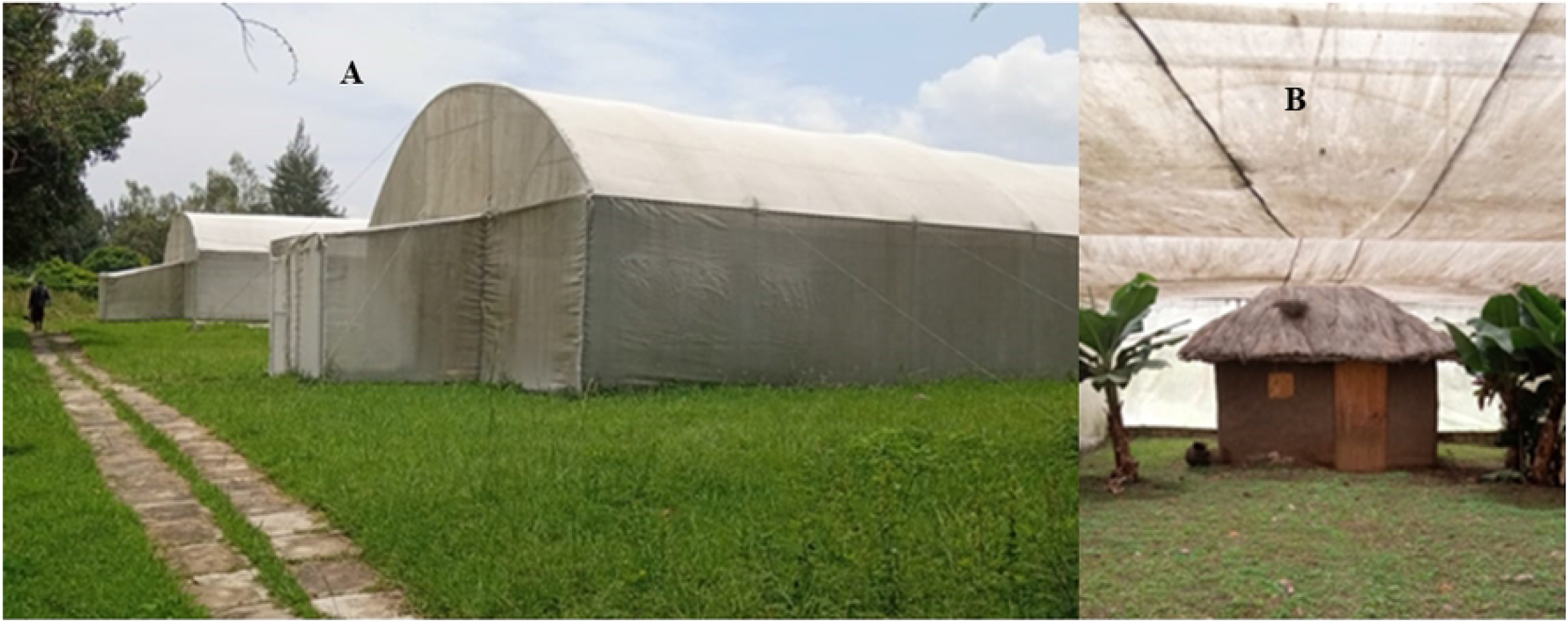
Pictures of (A), semi-field structures at KEMRI-CGHR, Kisumu and (B) a typical unmodified hut inside the semi-field structure

**Figure 2:**
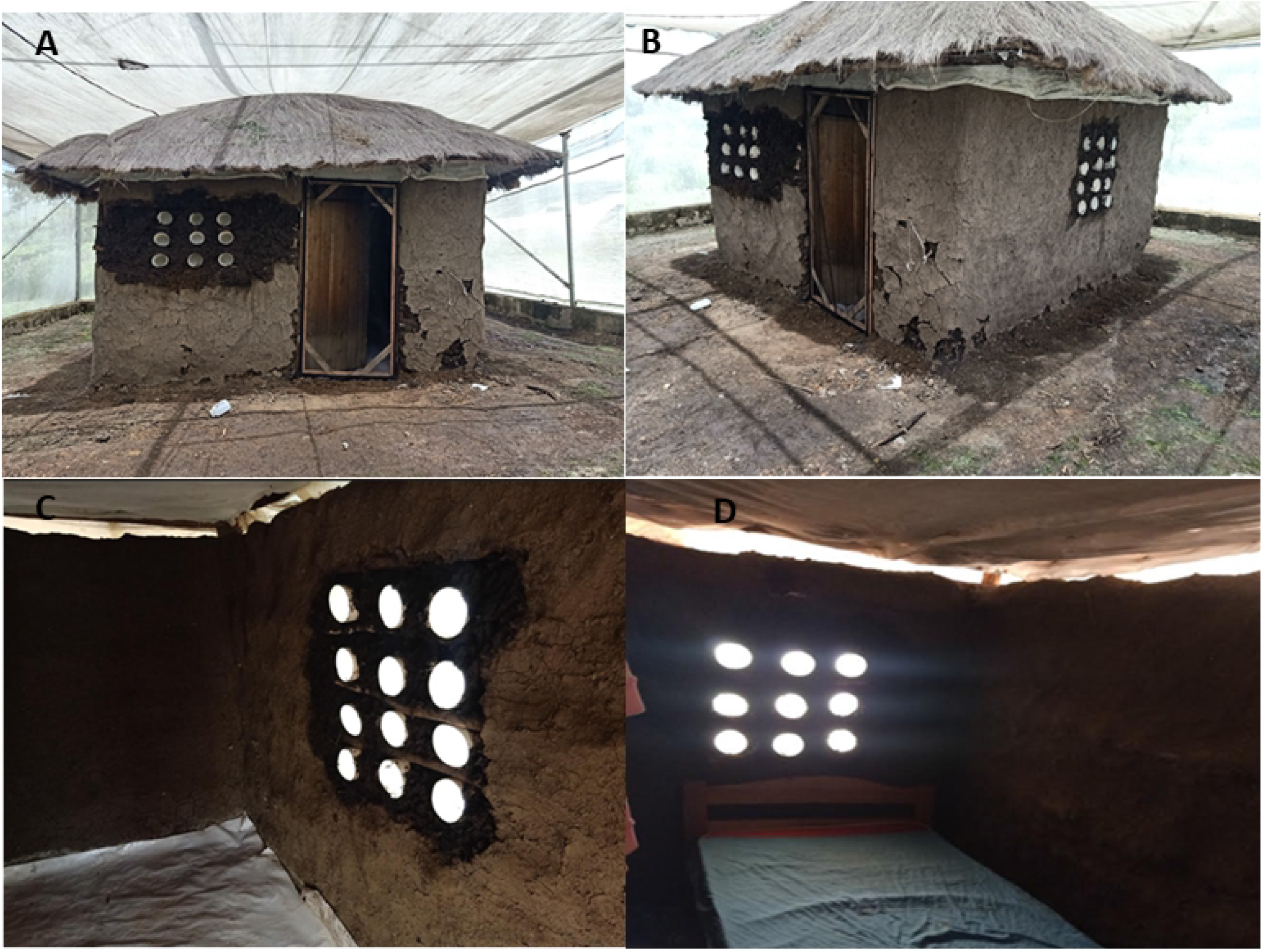
Picture of modified huts showing front and side views with air cavities and screened door (A and B), interior views of the living room C and bedroom (D) areas.

### Modification of the structures

Structural modifications were performed by replacing windows with screened air cavities and screening doors and eaves. The modifications were performed only for huts in Screen houses number 2 and 3 while screen houses number 1 and 4 were used as controls. The air cavities were made of PVC pipes of 200mm diameter and 150 mm length. The air cavities were fitted with insect mesh to allow air and light while blocking passage of mosquitoes. The air cavities were arranged into a panel of nine or twelve depending on the size of the window space. The gaps between the pipes were filled with mud, the same material used on the wall. Eaves were screened by attaching an insect mesh to the edge of the wall on one side and to the rafters and purlins on the roof on the other end to cover the eave space. The door to each hut was screened by introducing an openable wood framed screen shutter to the outside of the main door. An insect screen mesh was attached to the wood framed shutter using Velcro (Figure 2). For experiment 1, untreated insect screen material was used to screen the eaves whereas, in experiment 2, the insect mesh was treated with pirimiphos methyl.

### Mosquito Collections

Wild *Anopheles arabiensis* larvae were collected from rice fields in Ahero, Kisumu County and reared to adulthood at the KEMRI-CGHR insectaries in Kisumu. These were raised to 3-day old adults for the release experiments. Adult wild *An. funestus* were collected from Uranga in Siaya County and transported to KEMRI-CGHR insectaries for rearing of F1 generation. The gravid female *An. funestus* were provided with a laying pad inside a cage to collect eggs. Once eggs hatched, the resulting larvae were reared in the insectary conditions (27±2°C, 80±10% Humidity) to 3-day old adults for the release experiments. Laboratory reared, 3-day old adult, *An. arabiensis* Dongola strain were included in the release experiments. *Anopheles funestus* have high intensity (10X), resistance to pyrethroids while *An. arabiensis* from Ahero has low intensity (2X) resistance to the same class of insecticides. *Anopheles arabiensis* Dongola colony are a lab susceptible strain susceptible to all insecticides. All the mosquito populations were fully susceptible to organophosphates.

### Semi-Field Experiments

Two experiments were conducted. In experiment 1, the eaves of the modified huts were screened with untreated insect mesh material. In experiment 1, 5 releases of *An. arabiensis* from Ahero, 2 releases of *An. arabiensis* Dongola strain and 4 releases of *An. funestus* F1 generation from Siaya were done in the semi field structures over a total of 11 nights in March 2022.

In experiment 2, the eave screen was treated with Actellic® 300CS (pirimiphos-methyl). For this trial, 833ml of Actellic® 300CS was diluted in 6667mls litres of clean water in the 15-litre capacity H. D. Hudson Manufacturing Company (Chicago, IL) 67422 AD, Hudson X-pert spray pumps recommended by the WHO for use in IRS. The mixture was pressurized to 55 Psi [33]. The application rate was 1 m in 2.2s for 2m^2^ area of [34]. The netting material (length 4m by 1m width) was attached to a board, sprayed and allowed to dry under a shade following WHO guidelines [34]. It was wrapped in airtight polythene and kept at room temperature. In experiment 2, 5 releases of each of the three mosquito populations were performed over 15 nights in June 2022. Each release comprised 200 female mosquitoes per semi-field structure.

During each experiment, an adult male volunteer was recruited and slept under an untreated net in each of the huts. The volunteers were required to stay inside the huts from 2000HRS until 0600HRS the following morning except for bathroom breaks. Mosquitoes were released between 1800-1900hrs each evening from the middle of each semi field structure. Collection of the released mosquitoes was done in two sets, the first collections between 0600-0700HRS and a second and final collection between 0900-1000HRS the next day. Mosquito collection was done using mouth aspiration as well as mechanical aspiration using Prokopack aspirators. Mosquitoes collected indoors and those collected outdoors were kept in separate cups and labelled by the semi field structure number as well as the location of capture for counting.

### Data management and analysis

Recaptured mosquitoes were counted, and numbers recorded by trapping location as either indoors or outdoors of each hut for each mosquito population. Data were entered into Excel and imported into R statistical software version 4.1.2 for analysis. The risk ratio (RR) was used to assess the statistical significance of differences in mosquito densities between screened and unscreened huts. The over-dispersed data were fitted using Generalized Linear Mixed Model using Template Model Builder (glmmTMB)’ with a negative binomial distribution for the analysis of mosquito numbers between screened and unscreened huts. The total number of each *Anopheles* population recaptured indoors and outdoors of each hut were assessed as a function of intervention status (screened or unscreened) as a fixed effect while hut and sleeper being the random effect. To obtain the risk ratios (RR) and confidence intervals, we exponentiated the model coefficients.

## Results

Overall, the released mosquitoes were recaptured either inside or outside the huts within each semi field structure. In both experiment 1 and 2, the highest numbers of mosquitoes recaptured indoors were from unscreened huts compared to modified ones. While the numbers of mosquitoes of all species were relatively higher outdoors in semi field structures with screened huts compared to those with unscreened huts. The recapture rates were highest in *An. arabiensis* Dongola strain, followed by wild *An. funestus* from Siaya and last in wild. *An. arabiensis* from Ahero in both experiments 1 and 2 (Table 1).

**Table 1:** Numbers of mosquitoes recaptured indoors and outdoors of screened and unscreened huts within the semi field structures.

| Experiment | Species Strain | Treatment | Indoor | Outdoor | Number released | Number recaptured | % Recaptured |
| --- | --- | --- | --- | --- | --- | --- | --- |
| Experiment 1 | Ahero (*An. arabiensis*) | Hut1_ Unscreened | 109 | 85 | 1000 | 194 | 19.40 |
|  |  | Hut2_ Screened | 17 | 191 | 1000 | 208 | 20.80 |
|  |  | Hut3_ Screened | 1 | 185 | 1000 | 186 | 18.60 |
|  |  | Hut4_ Unscreened | 35 | 156 | 1000 | 191 | 19.10 |
|  | Dongola (*An. arabiensis*) | Hut1_ Unscreened | 120 | 53 | 400 | 173 | 43.25 |
|  |  | Hut2_ Screened | 15 | 213 | 400 | 228 | 57.00 |
|  |  | Hut3_ Screened | 6 | 231 | 400 | 237 | 59.25 |
|  |  | Hut4_ Unscreened | 80 | 188 | 400 | 268 | 67.00 |
|  | Siaya (*An. funestus*) | Hut1_ Unscreened | 117 | 93 | 800 | 210 | 26.25 |
|  |  | Hut2_ Screened | 18 | 209 | 800 | 227 | 28.38 |
|  |  | Hut3_ Screened | 8 | 229 | 800 | 237 | 29.63 |
|  |  | Hut4_ Unscreened | 135 | 85 | 800 | 220 | 27.50 |
| Experiment 2 | Ahero (*An. arabiensis*) | Hut1_ Unscreened | 140 | 165 | 1000 | 305 | 30.50 |
|  |  | Hut2_ Screened | 1 | 139 | 1000 | 140 | 14.00 |
|  |  | Hut3_ Screened | 0 | 150 | 1000 | 150 | 15.00 |
|  |  | Hut4_ Unscreened | 188 | 144 | 1000 | 332 | 33.20 |
|  | Dongola (*An. arabiensis*) | Hut1_ Unscreened | 354 | 124 | 1000 | 478 | 47.80 |
|  |  | Hut2_ Screened | 6 | 295 | 1000 | 301 | 30.10 |
|  |  | Hut3_ Screened | 12 | 333 | 1000 | 345 | 34.50 |
|  |  | Hut4_ Unscreened | 294 | 158 | 1000 | 452 | 45.20 |
|  | Siaya (*An. funestus*) | Hut1_ Unscreened | 147 | 140 | 1000 | 287 | 28.70 |
|  |  | Hut2_ Screened | 0 | 183 | 1000 | 183 | 18.30 |
|  |  | Hut3_ Screened | 0 | 204 | 1000 | 204 | 20.40 |
|  |  | Hut4_ Unscreened | 133 | 188 | 1000 | 321 | 32.10 |
| Total |  |  | 1936 | 4141 | 20800 | 6077 | 29.22 |

### Experiment 1

The mean number of each mosquito population recaptured indoors and outdoors of screened and unscreened huts within the semi field structure are presented in Figure 3. Relatively higher numbers of *An. arabiensis* from Ahero, *An. arabiensis* Dongola strain and *An. funestus* from Siaya were observed outdoors of screened Huts 2 and 3 compared to the numbers indoors. For unscreened huts, no difference in indoors and outdoors numbers of *An. arabiensis* from Ahero were observed in hut 1, except for hut 4 where relatively more were collected outdoors. Higher numbers of *An. arabiensis* Dongola strain were recaptured indoor in unscreened Hut 1, whereas, higher numbers were recaptured outdoors of hut 4. The numbers of *An. funestus* from Siaya were slightly higher indoor compared to outdoor, but the differences were not significant (Figure 3).

**Figure 3.**
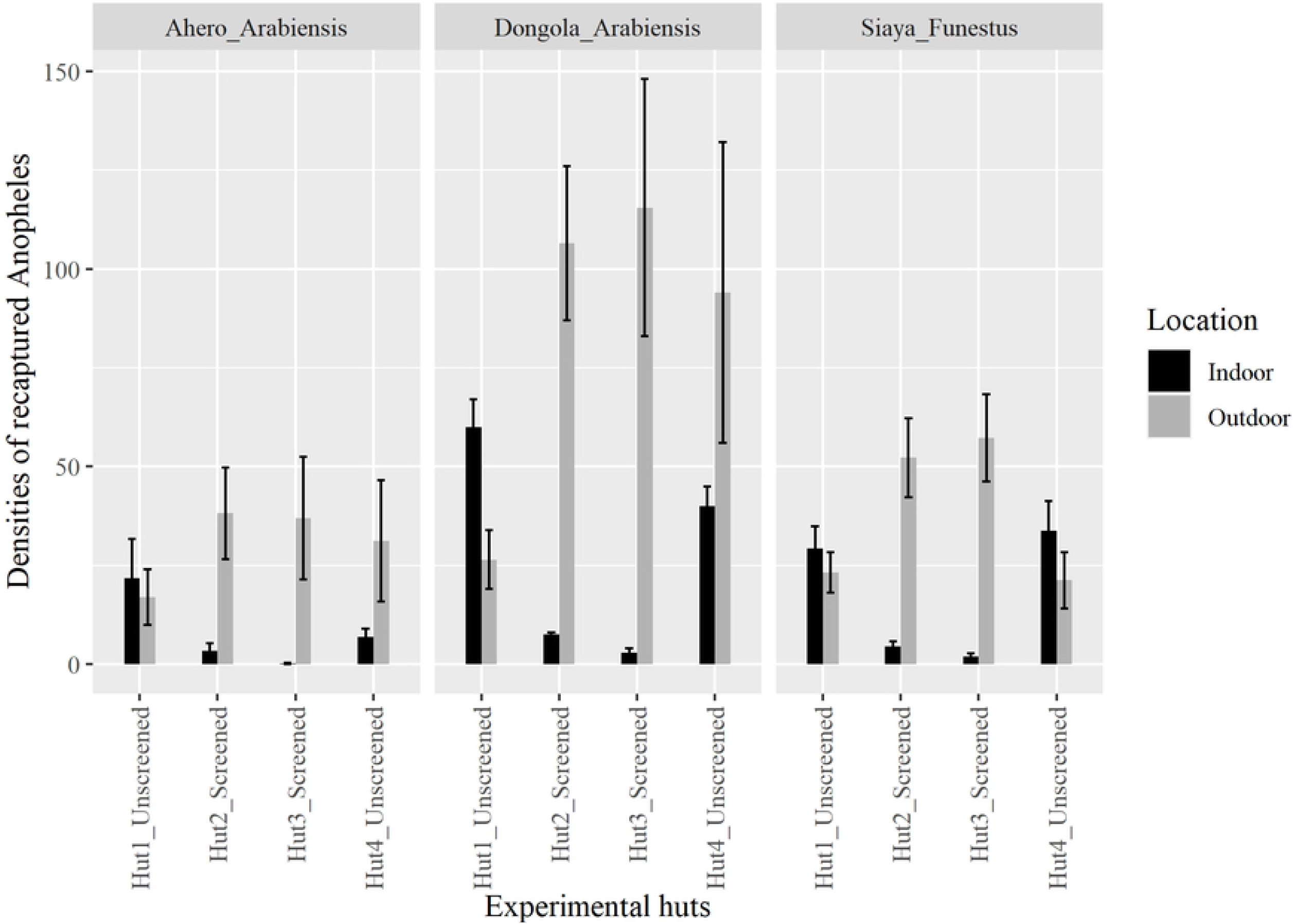
Mean number of *An. arabiensis* from Ahero, *An. arabiensis* Dongola strain and *An. funestus* from Siaya recaptured indoor and outdoor of screened and unscreened huts within the semi-field structures. Huts 1 and 4 were unscreened while Huts 2 and 3 were screened.

### Experiment 2

Relatively higher numbers of all the mosquito strains tested were recaptured outdoors compared to indoors for all screened Huts 2 and 3. No difference in numbers of *An. arabiensis* from Ahero were observed between indoor and outdoor of unscreened Huts 1 and 4. For *An. arabiensis* Dongola Strain, relatively more were recaptured indoors compared to outdoors of unscreened Huts 1 and 4. The numbers of *An. funestus* from Siaya were not different between indoors and outdoors of the unscreened Huts 1 and 4 (Figure 4).

**Figure 4.**
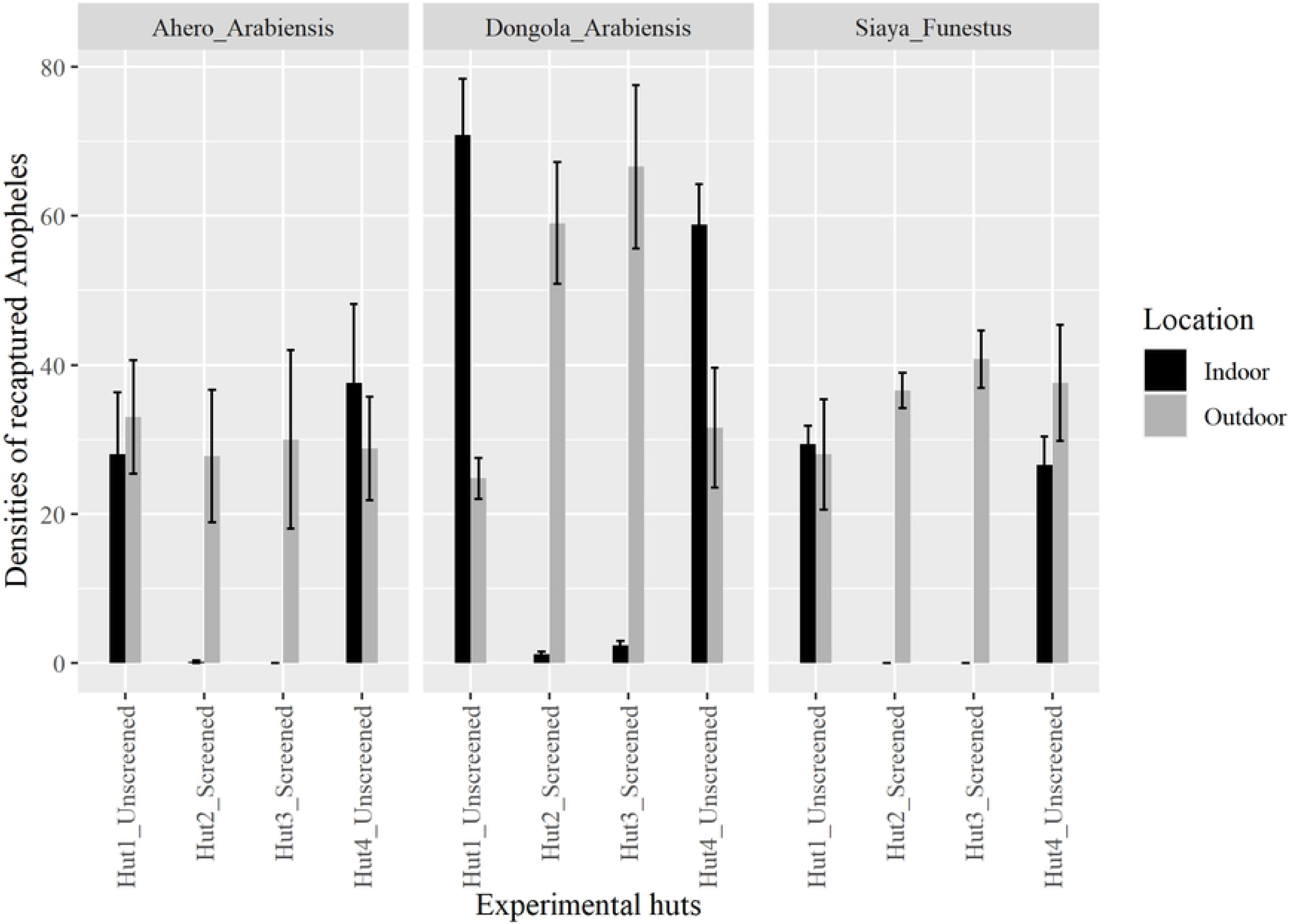
Mean number of *An. arabiensis* from Ahero, *An. arabiensis* Dongola strain and *An. funestus* from Siaya recaptured indoor and outdoor of screened and unscreened huts within the semi-field structures. Huts 1 and 4 were unscreened while Huts 2 and 3 were screened and the eave screens were treated with Actellic® 300CS (pirimiphos-methyl).

In experiment 1, significantly fewer *An. arabiensis* from Ahero [RR=0.10; (95%CI: 0.02-0.63); P=0.0145], *An. arabiensis* Dongola strain [RR=0.11; (95%CI: 0.04 – 0.19); P<0.0001 and *An. funestus* from Siaya [RR=0.10; (95%CI: 0.06-0.17); P<0.0001] were observed inside modified huts compared to unmodified ones. For outdoor collections, no significant difference was observed in the numbers of wild *An. arabiensis* from Ahero [RR=1.56; (95%CI: 0.72-3.38); P=0.259] and *An. arabiensis* Dongola strain [RR=2.10; (95%CI: 0.87-5.05); P=0.0985] between modified and unmodified huts. However, there were significantly more *An. funestus* from Siaya outside modified huts compared to unmodified ones [RR=2.46; (95%CI: 1.61-3.77); P<0.0001] (Table 2).

**Table 2:**
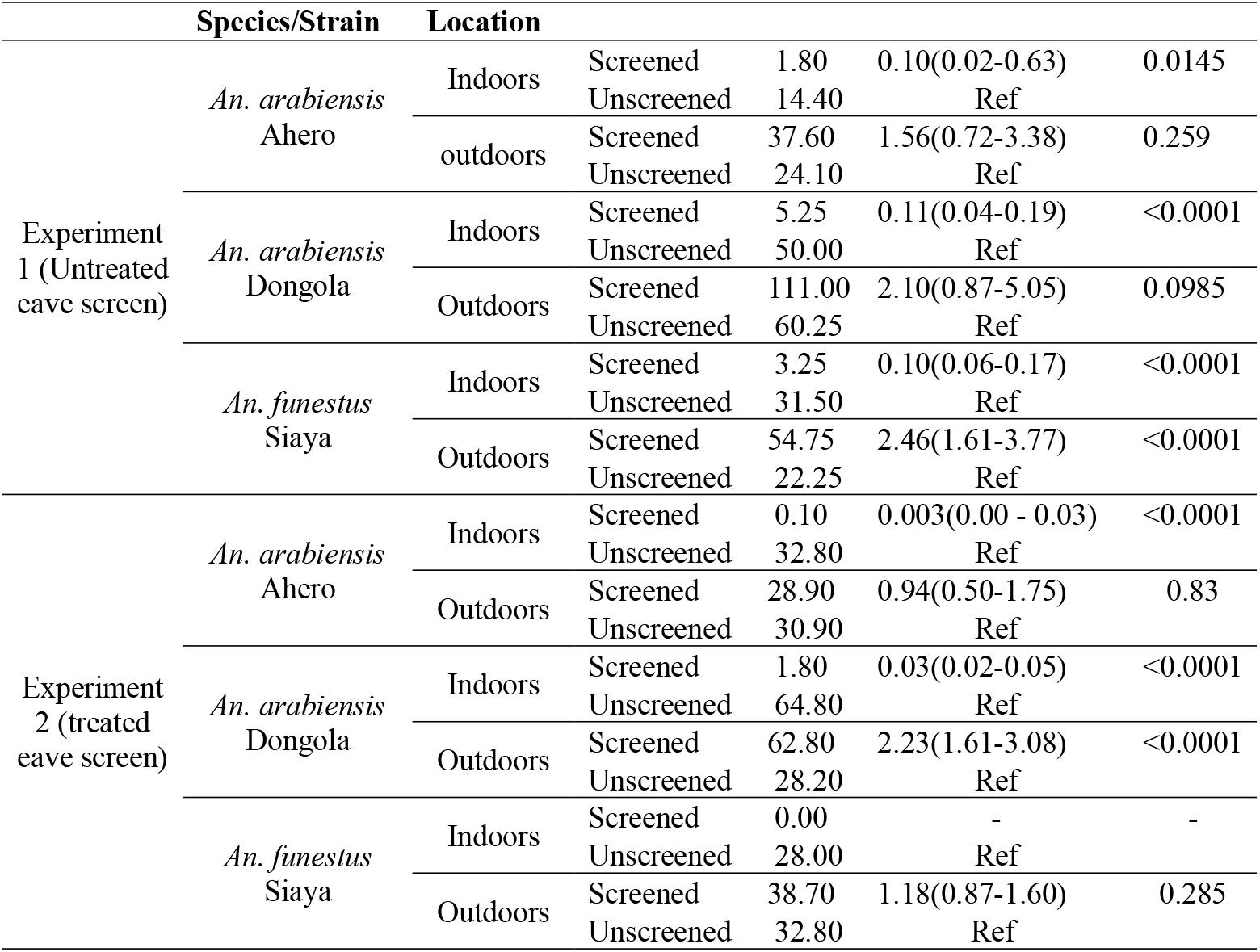
Comparison of mean *An. Arabiensis* from Ahero, *An. arabiensis* Dongola strain and *An. funestus* from Siaya between screened and unscreened huts within semi field structures. Experiment 1 involved eave screening with an untreated insect mesh while Experiment 2 used insect mesh treated with Actellic 300CS.

In experiment 2, huts with treated eave screen material had significantly reduced risk of *An. arabiensis* from Ahero [RR=00; (95%CI: 0.00-0.03); P<0.0001] and *An. arabiensis* Dongola strain [RR=0.03; (95%CI:0.02-0.05); P<0.0001] indoors compared unscreened ones. Hut screening with treated eave material eliminated the occurrence of *An. funestus* indoors. Significantly higher numbers of *An. arabiensis* Dongola strain were recaptured outdoors of screened huts compared to unscreened ones [RR=2.23; (95%CI: 1.61-3.08); P<0.0001]. No significant differences were observed for *An. arabiensis* from Ahero and *An. funestus* from Siaya outdoors in screened compared to unscreened huts (Table 2).

The densities of *An. arabiensis* from Ahero, *An. arabiensis* Dongola strain and *An. funestus* from Siaya recaptured inside and outside screened huts with untreated eave screen (Experiment 1) were compared to screened huts with treated eave screen (Table 3). Significantly fewer *An. arabiensis* from Ahero [RR=0.05; (95%CI: 0.00-0.77); P=0.0311] and *An. arabiensis* Dongola strain [RR=0.34; (95%CI: 0.18-0.64); P=0.000861] were recaptured indoors of huts with treated eave screen compared to huts with untreated eave screens. No *An. funestus* were recaptured indoor of huts with treated eave screens. In outdoor collections, significantly fewer *An. arabiensis* Dongola strain [RR=0.57; (95%CI: 0.41-0.78); P=0.000615] and *An. funestus* from Siaya [RR=0.71; (95%CI: 0.55-0.91); P=0.00607] were recaptured outside huts with treated eave screens compared to huts with untreated eave screens.

**Table 3:**
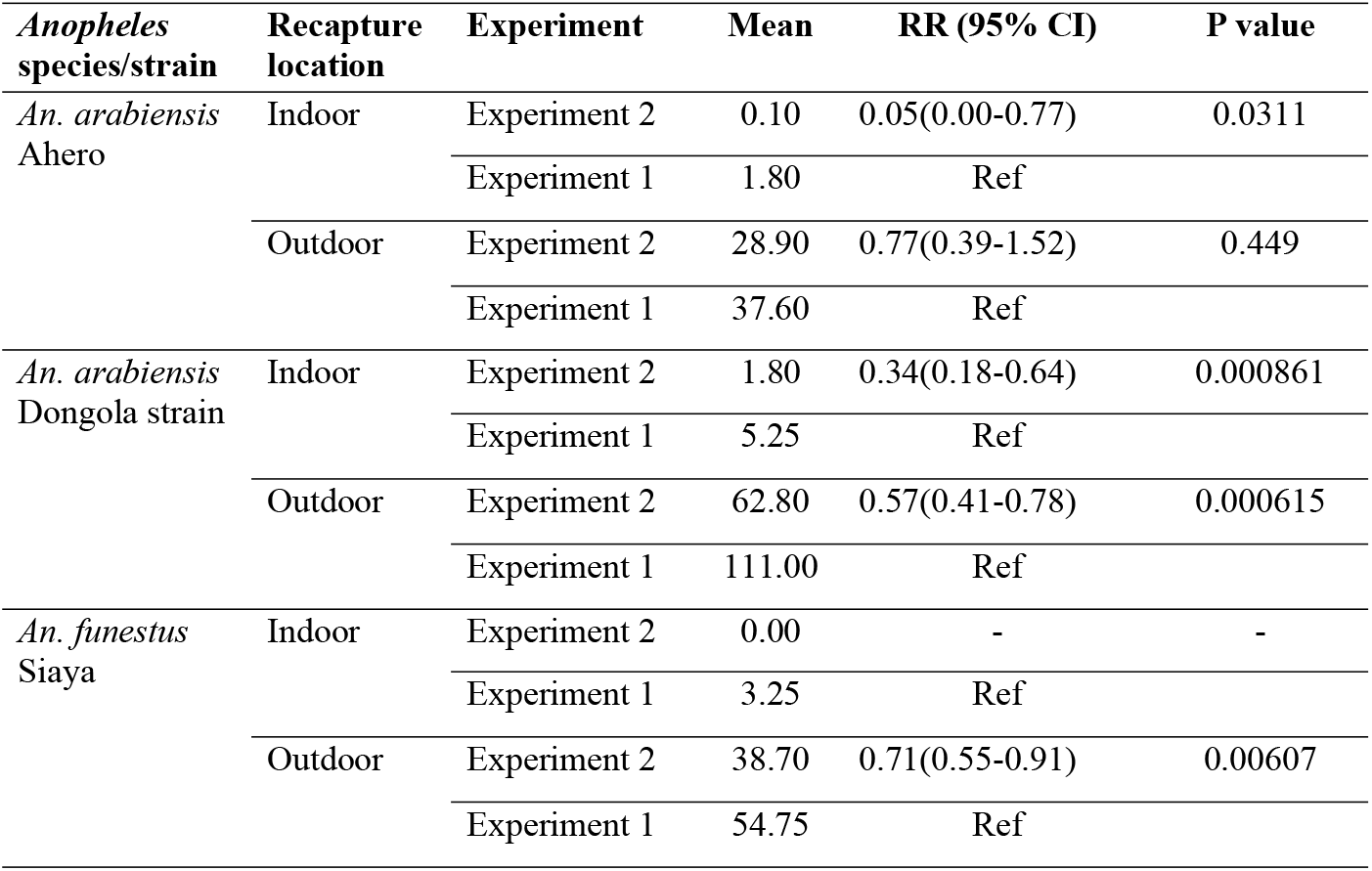
Comparison of mean number of *An. arabiensis* from Ahero, *An. arabiensis* Dongola strain and *An. funestus* from Siaya recaptured indoors and outdoors between screened huts with untreated eave screen (Experiment 1) and screened hut with treated eave screen (Experiment 2). Screen material used in Experiment 2 was treated with pirimiphos-methyl (Actellic 300CS).

## Discussion

Screening of doors, windows, and eaves significantly reduced indoor entry of *Anopheles* mosquitoes into huts under semi-field conditions. Treating the eave screen material with Actellic 300CS was observed to further reduce the numbers of *An. arabiensis* from Ahero and Dongola strains inside screened huts while eliminating the occurrence of *An. funestus* indoors. Additionally, screening was observed to prevent entry into the huts thus the increase of numbers of mosquitoes recaptured outside the huts, whereas, treating of eave screen material significantly reduced the numbers of *An. arabiensis* Dongola strain and *An. funestus* outdoors, within the semi field structures.

House screening has demonstrated potential for control of disease transmitting vectors such as mosquitoes [26, 28, 35-37]. Mosquitoes are adapted to enter and feed within human houses [18], hence transmitting malaria. Eaves are the main avenues for indoor entry of mosquitoes [18, 38], with doors and windows being additional routes. Blocking of eaves have been demonstrated to significantly reduce the numbers of mosquitoes indoors [28, 39] while additional screening of doors and windows increase the success of reducing mosquitoes indoors [35, 40]. In western Kenya, the major malaria vectors bite indoors, late and night [23-25] despite sustained use of ITNs. House screening therefore offers practical option for mosquito control in the region. Consistent with previous studies in the region, we observed inhibition of entry of local vector species *An. arabiensis* from Ahero and *An. funestus* from Siaya into huts under semi-field conditions.

Screening of huts was observed to limit entry of mosquitoes indoors while increasing the numbers recaptured outdoors within the semi-field structure. Similar observations are likely to be made in the actual field condition. Malaria vectors are usually closely connected with human dwellings [41], mostly occurring within the peri domestic space, either indoors or outdoors. One inadvertent result of house screening could be the increase in outdoor malaria transmission which remains poorly understood and controlled. This challenge has been previously observed in the implementation of ITNs, where increased coverage and use of ITNs has been associated with outdoor malaria transmission [14, 16, 17, 42, 43]. Insect proof housing is more likely to exacerbate the already challenging outdoor malaria transmission, therefore vector population reduction options need to be explored in combination with house screening.

Treating of screening material with an effective insecticide presents an option for vector population reduction. We observed treating of eave screen material to significantly reduce the numbers of *An. arabiensis* Dongola strain both indoors and outdoors, *An. arabiensis* Ahero strain indoor and *An. funestus* outdoors while eliminating the numbers indoors under semi field conditions. *Anopheles* mosquitoes mostly enter and exit houses through the eaves [18], therefore, treating of eave screens offers a viable option for reduction of mosquito populations. Consistent with these observations, eave tubes treated with pyrethroids have been demonstrated to be effective in preventing house entry and reducing mosquito population under same field conditions [44] and in the community [45]. Combination of house screening with delivery of insecticides on the eaves screen material offers greater protection against endophilic and endophagic mosquitoes. However, care should be taken in the choice of insecticides to ensure cost effectiveness, efficacy and mitigate against the spread of insecticide resistance in mosquitoes.

The different population of malaria vector species used in this evaluation were affected differently by house screening and treatment of eave screen material. Screening significantly reduced the numbers of *An. arabiensis* from Ahero indoors, however there was no significant difference in the numbers outdoors between screened and unscreened huts. Similar results were observed when eave screen was treated with Actellic 300 CS. Furthermore, the treatment of eave screens only reduced the numbers indoors but did not have an impact on the numbers outdoor. These observations are consistent with reported behaviour of *An. arabiensis* in western Kenya. The species has been reported to be more exophagic and exophilic [46, 47]. Consequently, the numbers observed in this study to have entered the unscreened huts were not many such that no difference was observed in the numbers outdoors between the two treatment groups.

*An. arabiensis* Dongola strain exhibited the highest recapture rate both indoors and outdoors compared to the field caught species. Screening significantly reduced indoor entry of the species and further reductions were observed with treatment of eave screen material. The response of *An. arabiensis* Dongola strain to house screening may not reflect the actual field situation since the species used is a laboratory colony.

House screening had significant reductions in the numbers of *An. funestus* indoor while increasing the numbers outdoors. Treating of eaves screen material with Actellic 300CS eliminated the species indoors while significantly reducing the numbers outdoor when compared with untreated eave screen material. *An. funestus* in western Kenya has been reported to be highly endophilic and endophagic [46, 47] hence feeding and resting more frequently indoors. The species is therefore more inclined to indoor entry. Consistent with these reports, we observed screened huts to have significantly lower numbers indoors compared to unscreened huts, whereas the numbers outdoor of screened huts were higher than those around unscreened huts. Treatment of eave screen material further affected the population of *An. funestus* indoors and outdoors of the huts. Previous studies in the region have reported the species to be highly susceptible to an effective insecticide. *An. funestus* was reduced to near elimination in the Asembo bay area with introduction of ITNs [48]. Similar results were observed in Migori County, western Kenya, following a single indoor residual spray (IRS) campaign with Actellic 300CS [25].

## Conclusion

Screening of huts significantly reduces the risk of *Anopheles* mosquitoes’ entry indoor under semi-field condition. Treatment of the eave screen material further reduced the vector populations indoors and outdoors with potential elimination of occurrence of *An. funestus* indoor. Consequently, an evaluation of insect proof housing, coupled with delivery of an effective insecticide on the eave screen material at a community level is recommended for demonstration of epidemiological impact.

## Declarations

### Ethics approval and consent to participate

The study was approved by the Kenya Medical Research Institute/ Scientific and Ethics Review Unit SERU 2776. Written consents were obtained from all study volunteers who slept in the huts.

### Consent for publication

Not applicable.

### Disclaimer

The findings and conclusions in this report do not necessarily reflect the official position of the U.S. Department of Health and Human Services or the U.S. Centers for Disease Control and Prevention. The use of trade names is for identification only and does not imply endorsement by the U.S. Department of Health and Human Services or the U.S. Centers for Disease Control and Prevention.

## Authors’ contributions

JO and JS conceived the intervention and managed the innovation process that led to house screening. BA and EO wrote the protocol. BA, SA and EO conducted the study. BA and VM performed the data analysis. BA wrote the manuscript; all authors reviewed and approved the manuscript.

## Data availability

Data available at request from the corresponding author.

## Acknowledgements

We would like to thank the study volunteers who slept in the experimental huts to attract mosquitoes and the KEMRI Entomology staff who took part in this evaluation.

